# Near-atomic resolution reconstructions from *in situ* revitrified cryo samples

**DOI:** 10.1101/2023.02.20.529238

**Authors:** Gabriele Bongiovanni, Oliver F. Harder, Jonathan M. Voss, Marcel Drabbels, Ulrich J. Lorenz

## Abstract

We have recently introduced a microsecond time-resolved version of cryo-electron microscopy (cryo-EM) to enable the observation of the fast conformational motions of proteins. Our technique involves locally melting a cryo sample with a laser beam to allow the proteins to undergo dynamics in liquid phase. When the laser is switched off, the sample cools within just a few microseconds and revitrifies, trapping particles in their transient configurations, in which they can subsequently be imaged. We have previously described two alternative implementations of the technique, using either an optical microscope or performing revitrification experiments *in situ*. Here, we show that it is possible to obtain near-atomic resolution reconstructions from *in situ* revitrified cryo samples. Moreover, the resulting map is indistinguishable from that obtained from a conventional sample within our spatial resolution. Interestingly, we observe that revitrification leads to a more homogeneous angular distribution of the particles, suggesting that revitrification may potentially be used to overcome issues of preferred particle orientation.

**Synopsis:** Near-atomic resolution reconstructions can be obtained from *in situ* melted and revitrified cryo samples. Revitrification results in a more homogeneous angular distribution.

## Introduction

Proteins play a crucial role in most biological processes. They catalyze reactions, regulate gene expression, receive and transmit biological signals, and participate in the recognition of pathogens (Benkovic and Hammes-Schiffer 2003; Buccitelli and Selbach 2020; Wootten et al. 2018; Kumar, Kawai, and Akira 2009). To perform their tasks, they undergo large-scale domain motions that bear resemblance to those of manmade machines (Alberts 1998). However, because of the fast timescale of these motions, typically microseconds, it has largely remained impossible to observe them directly, which has left our understanding of protein function fundamentally incomplete (Henzler-Wildman and Kern 2007; Otosu, Ishii, and Tahara 2015; Persson and Halle 2008). In fact, it has been argued that understanding protein function represents the next frontier in structural biology (Ourmazd, Moffat, and Lattman 2022).

We have recently introduced a novel approach to time-resolved cryo-EM that affords microsecond time resolution. It thus promises to significantly advance our understanding of protein function by enabling direct observations of the motions of proteins (Voss et al. 2021b; 2021a; Harder et al. 2022). Our approach involves rapidly melting a cryo sample with a laser beam. Once the sample is liquid, conformational dynamics are triggered with a suitable stimulus, for example by releasing a caged compound or directly exciting a photoactive protein with a second laser pulse (Klein-Seetharaman 2002; Ellis-Davies 2007; Shigeri, Tatsu, and Yumoto 2001). As the particle dynamics unfold, the heating laser is switched off at a well-defined point in time, and the sample revitrifies within just a few microseconds, trapping the particles in their transient configurations, in which they can subsequently be imaged with conventional cryo-EM techniques (Voss et al. 2021b).

We have previously described two different implementations of our technique. Melting and revitrification experiments can be performed in an optical microscope, using a correlative light-electron microscopy approach (Bongiovanni et al. 2022). Alternatively, such experiments can be carried out *in situ* (Voss et al. 2021a; 2021b; Harder et al. 2022) with a modified transmission electron microscope (Olshin, Drabbels, and Lorenz 2020). *In situ* experiments offer the advantage that electron micrographs of revitrified areas can provide an immediate feedback on conformational changes of the proteins. It is even possible to perform on-the-fly reconstructions to guide the search for suitable experimental parameters. Moreover, it is straightforward to determine the diameter of the revitrified area from electron micrographs, which can be monitored to adjust the laser power in revitrification experiments, as we have previously described (Voss et al. 2021a). A potential drawback of *in situ* experiments is that the sample has to be exposed to a small electron dose (about 10^-3^ electrons/Å^2^) in order to locate a suitable area and aim the laser beam. This is a potential issue since we have previously observed that exposure to a dose as low as a few electrons/Å^2^ prior to revitrification induces enough beam damage to cause the particles to disassemble once the sample has turned liquid (Voss et al. 2021b). We have recently demonstrated that *in situ* revitrification does not alter the structure of the proteins (Harder et al. 2022). However, the resolution of the reconstructions we obtained was limited by the performance of our instrument to a few Angstroms. This leaves open the question whether *in situ* revitrification might induce structural changes that only become evident at higher spatial resolution or might even limit the obtainable resolution.

Here, we show that near-atomic resolution reconstructions can be obtained from *in situ* revitrified cryo samples by transferring them to a high-resolution electron microscope for cryo imaging. Interestingly, our analysis reveals that rapid melting and revitrification reshuffles the particles, creating a more homogeneous angular distribution.

## Results and discussion

**Figure 1** illustrates the experimental workflow. A cryo sample of mouse heavy-chain apoferritin was prepared on a UltrAuFoil specimen support, which features a holey gold film (1.2 μm diameter holes) on a 300 mesh gold grid (**Figure 1a**). *In situ* melting and revitrification experiments were then performed with a JEOL 2200FS transmission electron microscope that we have modified for time-resolved experiments (**Figure 1b**) (Olshin, Drabbels, and Lorenz 2020). The melting laser (532 nm) is directed at the sample with the help of an aluminum mirror placed above the upper pole piece of the objective lens, so that the laser beam strikes the sample at close to normal incidence. To perform a melting and revitrification experiment, the laser beam was aimed at the center of a grid square, and a 20 μs laser pulse was used to revitrify an area of typically about 25 holes. The sample was then transferred to a Titan Krios G4 (Thermo Fisher Scientitic) for high-resolution imaging (**Figure 1c**). Micrographs were collected from 16 revitrified areas as well as 11 conventional areas that had not been exposed to the laser beam. Finally, single-particle reconstructions of both data sets were performed with CryoSPARC (Punjani et al. 2017) to yield near-atomic resolution maps (**Figure 1d**).

**Figure 1.**
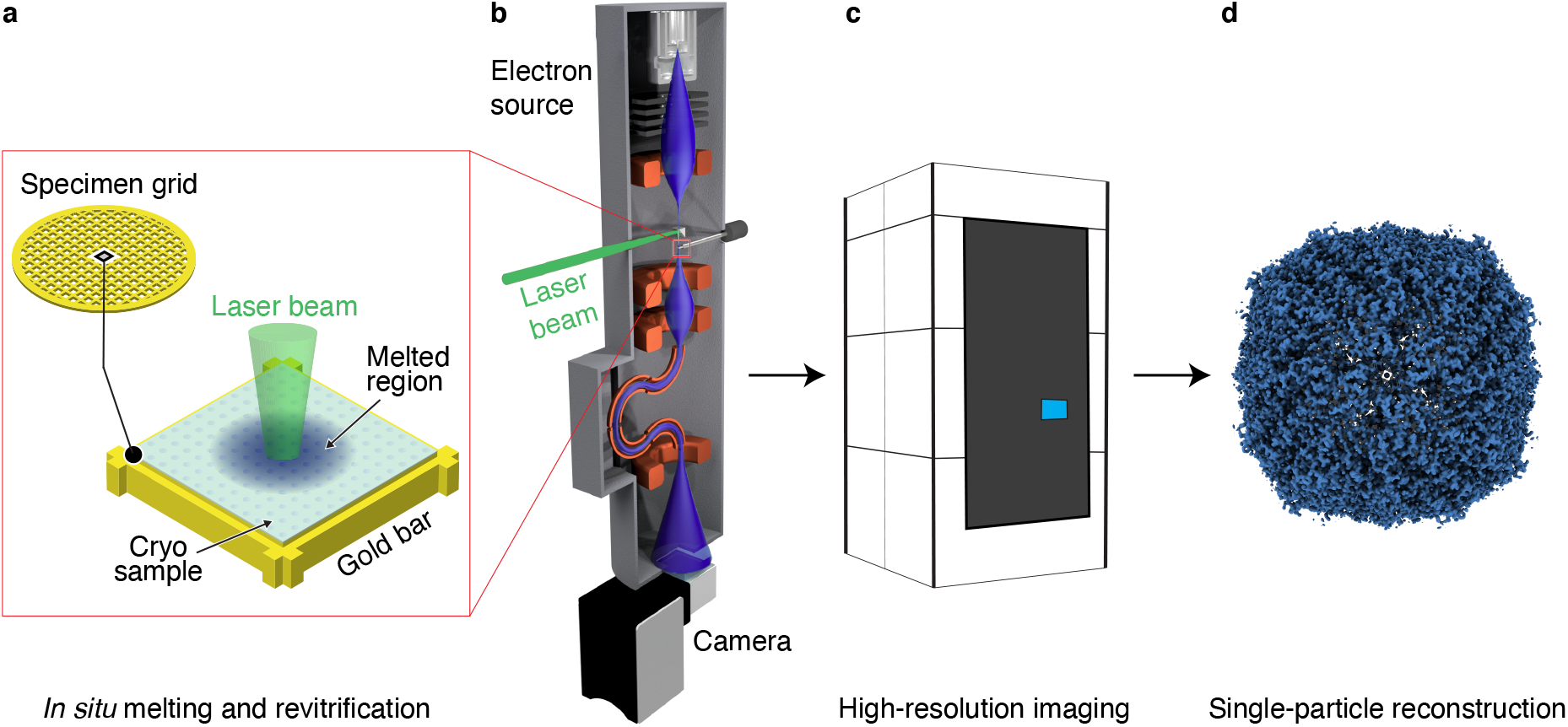
Illustration of the workflow to obtain for near-atomic resolution reconstruction from *in situ* revitrified cryo samples. (**a,b**) Cryo samples on UltrAuFoil grids are melted and revitirified *in situ* with a modified JEOL 2200F transmission electron microscopes. (**c**) The samples are then transferred to a Titan Krios G4 for high-resolution imaging. (**d**) Single-particle reconstructions from the revitrified areas yield near-atomic resolution maps.

**Figure 2** compares the reconstructions obtained from the conventional (left) and revitrified areas (right). Within our resolution (1.61 Å and 1.59 Å, respectively), the maps are undistinguishable. This is also evident in the details of side chains shown in **Figure 2b**, which reveal well-resolved densities of aromatic rings as well as individual water molecules. A model of mouse heavy-chain apoferritin is shown (Wu, Lander, and Herzik 2020) that we have placed into the density of through rigid-body fitting. These results confirm our previous observations that the *in situ* melting and revitrification process leaves the proteins intact (Harder et al. 2022). Moreover, we can determine to atomic-scale precision that under our experimental conditions, *in situ* revitrification does not alter the structure of the particles, nor does reduce the obtainable spatial resolution. Evidently, the small electron dose of about 10^-3^ electrons/Å^2^ that the sample is exposed to in an *in situ* experiment prior to revitrification does not cause sufficient beam damage to deteriorate the resolution.

**Figure 2.**
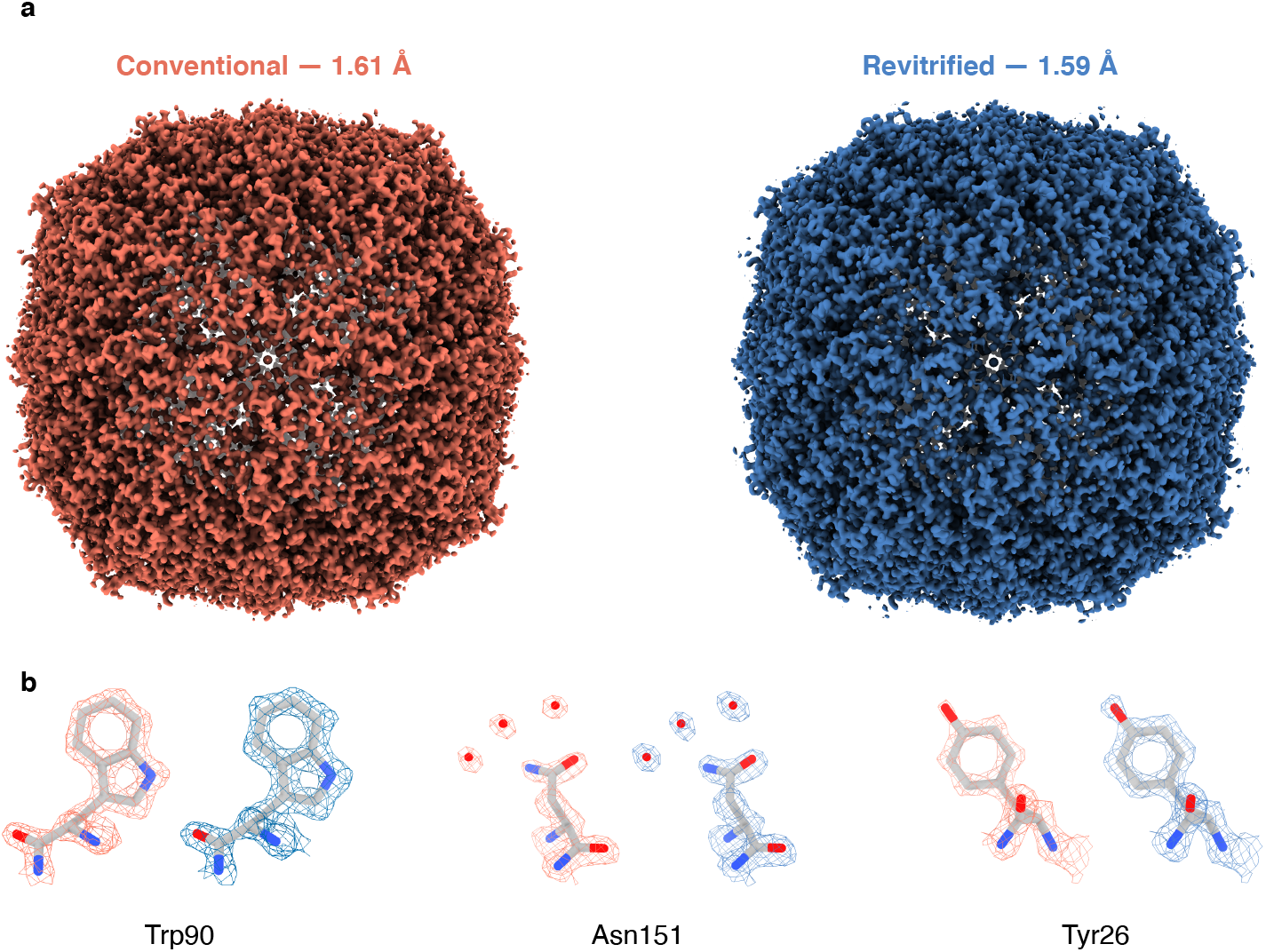
Near-atomic resolution reconstructions of apoferritin from conventional and revitrified areas. (**a**) The maps from conventional (left) and revitrified areas (right) are indistinguishable within the resolution obtained (~1.6 Å). (**b**) Details of the reconstructions show well-resolved side chain densities as well as individual water molecules. A model of apoferritin (Wu, Lander, and Herzik 2020) is placed into the density through rigid body fitting.

Interestingly, melting and revitrification results in a more homogeneous angular distribution of the particles. This is evident in **Figure 3a**, which compares the distributions for the conventional (left) and revitrified sample areas (right). Preferred particle orientations in cryo samples are thought to arise due to the adhesion of hydrophobic regions of the protein surface to the air-water interface (Taylor and Glaeser 2008). Such effects are small for apoferritin, which exhibits little orientational preference. The angular distribution features six narrow, symmetry equivalent maxima. Each is surrounded by eight broad, shallow minima, which fall into two groups of four that are related to each other by symmetry. After revitrification, this distribution is markedly broadened, with the ratio of the populations of the most and least probable orientation decreasing from 1.6 to 1.3.

**Figure 3.**
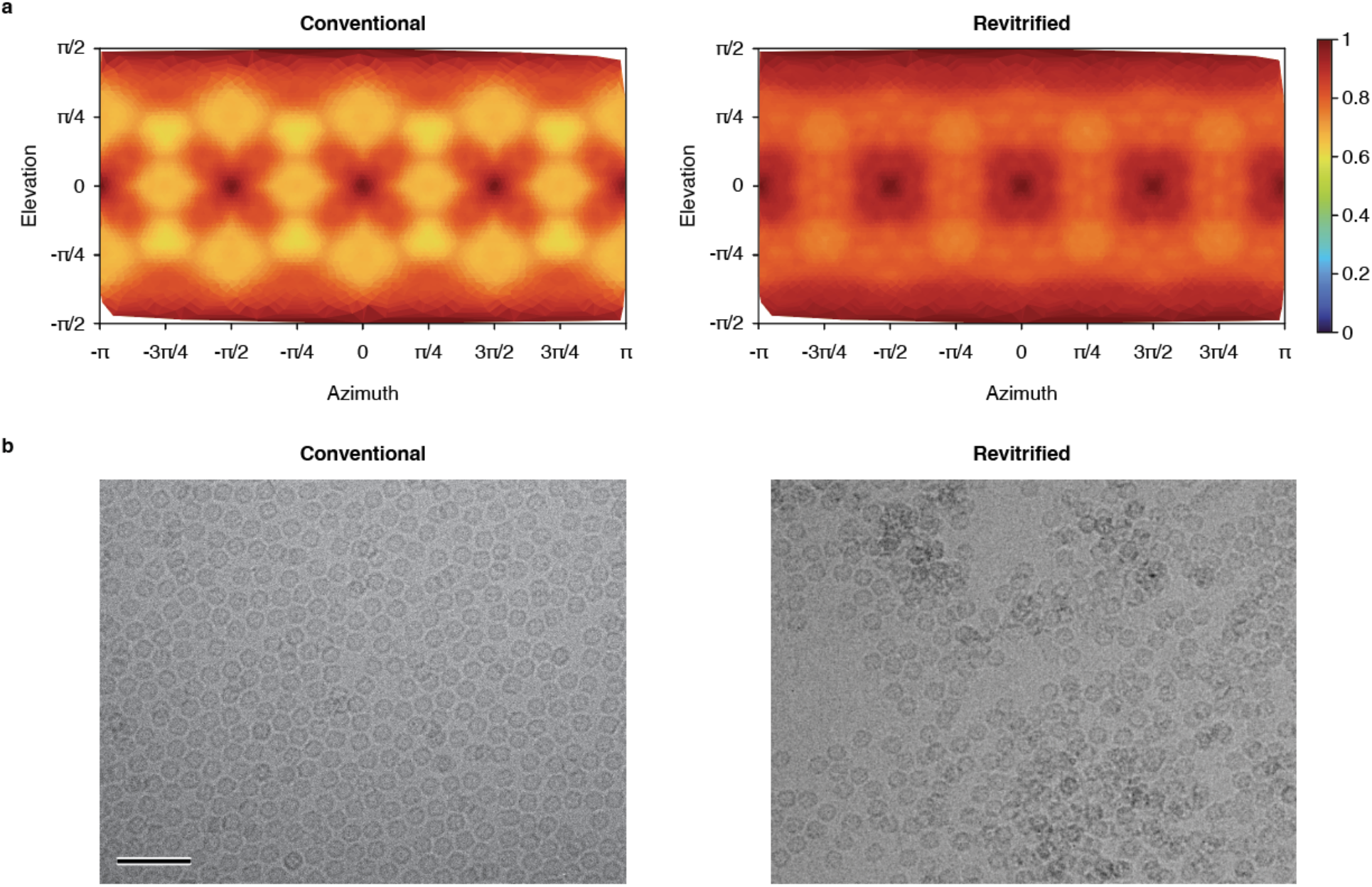
Revitrification reshuffles the particles, which results in a more homogeneous angular distribution. (**a**) Distribution of particle views in the conventional (left) and revitrified sample areas (right). The distributions are shown with octahedral symmetry applied. (**b**) A micrograph of a conventional sample area shows a homogeneous distribution of the particles (left), wheras an uneven particle distribution is observed in revitrified areas (right).

The broadening of the angular distribution upon revitrification is accompanied by changes in the spatial distribution of the particles, which provides hints as to the underlying mechanism. **Figure 3b** shows that while conventional samples typically feature a homogeneous particle distribution (left), revitrification results in an uneven distribution, with the particles clustering in some areas. Evidently, rapid laser melting of the sample reshuffles the particles and creates a non-equilibrium spatial and angular distribution. The time window during which the sample is liquid (< 20 μs) is manifestly insufficient to reestablish an equilibrium distribution as it is initially prepared during the plunge freezing process, where the sample is given about 1 s to equilibrate between blotting and plunging. Revitrification therefore traps a non-equilibrium distribution. This is also consistent with the expected rotational timescales. The rms rotation angle of freely diffusing apoferritin is about 80 degrees on a timescale of 10 μs. However, near a surface, particles frequently experience so-called anomalous diffusion, with diffusion coefficients lowered by several orders of magnitude (Jamali et al. 2021). The rms rotation angle in our experiment is therefore likely lower than the separation between the maxima and minima of the angular distribution (**Figure 3a**).

Several phenomena may potentially explain the reshuffling of the particles during the melting process. As we have previously described, the sample partially crystallizes during rapid laser heating when it traverses so-called no man’s land (Voss et al. 2021a). It is conceivable that the formation of ice crystallites and their subsequent melting exert small forces upon the particles that scramble their distribution. Another possibility is that stresses present in the vitreous ice film that build up during plunge freezing due to the difference in expansion coefficient between gold and water (Thorne 2020) are unevenly released during rapid laser melting, causing a microscopic flow of the liquid to reshuffle the particles. Further experiments will be needed to definitively settle this point.

## Conclusion

Our experiments demonstrate that near-atomic resolution reconstructions can be obtained from *in situ* melted and revitrified cryo samples, in agreement with previous results for samples that we revitrified in an optical microscope (Bongiovanni et al. 2022). This strongly suggests that the revitrification process does not impose a fundamental limit on the obtainable spatial resolution. Evidently, the small electron dose that the sample is exposed to in an *in situ* experiment prior to melting and revitrification does not alter the protein structure or the achievable resolution. This is an important result since *in situ* experiments offer the advantage over revitrification with an optical microscope that they can provide an immediate feedback, which speeds up the search for suitable experimental parameters.

The observed scrambling of the angular distribution of the particles during revitrification may potentially offer a means to overcome issues with preferred orientation that in some samples cause views to be underrepresented or even absent, making it difficult to obtain a reconstruction (Tan et al. 2017). While tilting the sample during imaging can be used to mitigate this effect (Penczek and Frank 2006; Tan et al. 2017), tilting to high angles reduces the obtainable resolution (Tan et al. 2017). Instead, laser revitrification may potentially be employed to repopulate hidden views without the need for tilting. Further experiments will be needed to determine whether this will be possible in samples with a strong preferred orientation and to find experimental parameters that maximize the effect.

## Methods

Cryo specimens were prepared on UltrAuFoil R1.2/1.3 300 mesh grids (Quantifoil). The grids were plasma cleaned for 1 min to render them hydrophilic (Tedpella “EasyGlow”). Subsequently, 3 μl of the sample solution (mouse heavy chain apoferritin, 8.5 mg/ml in 20 mM HEPES buffer with 300 mM sodium chloride at pH 7.5) were applied, and the samples were plunge-frozen using a Vitrobot Mark IV (Thermo Fisher Scientific, 3 s blotting time, 95% relative humidity at a temperature of 10 °C).

*In-situ* revitrification experiments were carried out with a modified JEOL 2200FS transmission electron microscope (Olshin, Drabbels, and Lorenz 2020), using a Gatan Elsa single-tilt cryo holder. Microsecond laser pulses for melting and revitrification (76 mW) were obtained by modulating the output of a 532 nm continuous wave laser (Laser Quantum, Ventus 532) with an acousto-optic modulator (AA optoelectronics). The beam was focused in the sample plane, giving an elliptical spot of 62 μm × 165 μm FWHM, as determined from a knife edge scan. Before transferring the sample for high-resolution imaging, an atlas was recorded, which allowed us to readily identify the revitrified areas later.

The micrographs in **Figure 3b** were recorded with the JEOL 2200FS. High-resolution micrographs for single-particle reconstructions (**Figure 2** and **3**) were collected on a Titan Krios G4 (Thermo Fisher Scientific), which was operated at 300 kV acceleration voltage. Zero-loss filtered images (Selectris X energy filter, 10 eV slit width) were recorded with a Falcon 4 camera, using an exposure time of about 2.5 s for a total dose of 50 electrons/Å^2^. The pixel size was set to 0.455 Å. Defocus values were in the range of 0.5–1.2 μm.

Single-particle reconstructions were performed in cryoSPARC 3.3.1 (Punjani et al. 2017) (details in the Supporting Information). The conventional (revitrified) dataset — both recorded on the same specimen grid — contained 4745 (1591) images, which were patch motion corrected. After CTF estimation, 1815 (1552) micrographs with a resolution better than 6 Å were retained. Template-based particle picking yielded 114414 (88004) particles. After two rounds of 2D classification, 91541 (74468) particles were retained, which were used for *ab initio* reconstruction (*C1* symmetry), followed by heterogeneous refinement using 2 classes (*C1* symmetry). The 72270 (72811) particles in the most populated class were then used for homogeneous refinement with *O* symmetry, giving a map with 1.61 Å (1.59 Å) resolution.

## Supporting information

Supplementary Information for the manuscript

## Associated Content

### Supplementary Material

Details of the single-particle reconstruction workflow Supplementary images

### Data Availability Statement

The data that support the findings of this study are available from the corresponding author upon reasonable request. The maps of apoferritin obtained from conventional and revitrified sample areas have been deposited on EMDB (EMD-15715 and EMD-15721, respectively), and the corresponding data sets on EMPIAR (EMPIAR-11345 and [EMPIAR code to be updated]).

### Author Information

**Corresponding author:** Ulrich J. Lorenz

**Notes:** The authors declare no competing financial interests.

## Acknowledgements

This work was supported by the ERC Starting Grant 759145 and by the Swiss National Science Foundation Grant PP00P2_163681. We acknowledge the support of the Dubochet Center for Imaging (DCI) in Lausanne. We also thank Dr. Kevin Lau (PTPSP EPFL, Switzerland) for providing the apoferritin samples.

