## Supplementary Information for the manuscript for "Near-atomic resolution reconstructions from *in situ* revitrified cryo samples"

**Supplementary Material 1. Single-particle reconstructions**

The conventional (revitrified) dataset was processed in CryoSPARC 3.3.1 (Punjani *et al.*, 2017), using 40 fractions an no upsampling. After patch motion correction, CTF estimation yielded 1815 (1552) images with a resolution better than 6.0 Å and relative ice thickness between 1.0 and 1.15, which were retained for further processing. For the revitrified data set, care was taken exclude images from outside of the revitrified areas.

Particle picking was conducted with the following procedure. Initially, we applied blob picking to the images in the conventional dataset, using a radius between 110 and 120 Å. We then selected 127113 of the 228751 particles picked, based on their NCC and power score, and extracted them with a box size of 560 pixels. Afterwards, we performed one round of 2D classification, with 50 classes. We selected the 16 classes of the highest quality, containing 53418 particles, and used them as a template for template-based particle picking on both datasets. This procedure yielded 730858 (672391) particles, which we selected based on their NCC and power scores. Finally, we extracted the 146702 (113632) remaining particles with a box size of 560 pixels, yielding a total of 114414 (88004) particles.

Reconstructions were obtained with the following procedure. The 114414 (88004) selected particles underwent a first round of 2D classification, with 50 classes. The 92890 (76655) particles belonging to the 12 (14) best classes underwent a second round of 2D classification, this time using a circular mask of 150 Å and 30 classes. The best 15 (14) classes, containing 91541 (74468) particles, were manually selected and used for ab-initio reconstruction, with 2 classes and *C1* symmetry. Particles belonging to both classes were then used for a round of heterogeneous refinement, using 2 classes and *C1* symmetry. The 90590 (72811) particles belonging to the most populated class were used for a final homogeneous refinement, imposing *O* symmetry. This last refinement step was performed using per-particle defocus optimization, per-group CTF refinement, and implementing Ewald sphere and anisotropic magnification correction. The procedure yielded a final map at 1.61 Å (1.59 Å), as determined by the Gold Standard Fourier Correlation Shell (GSFSC) at 0.143.

### Supplementary Figures

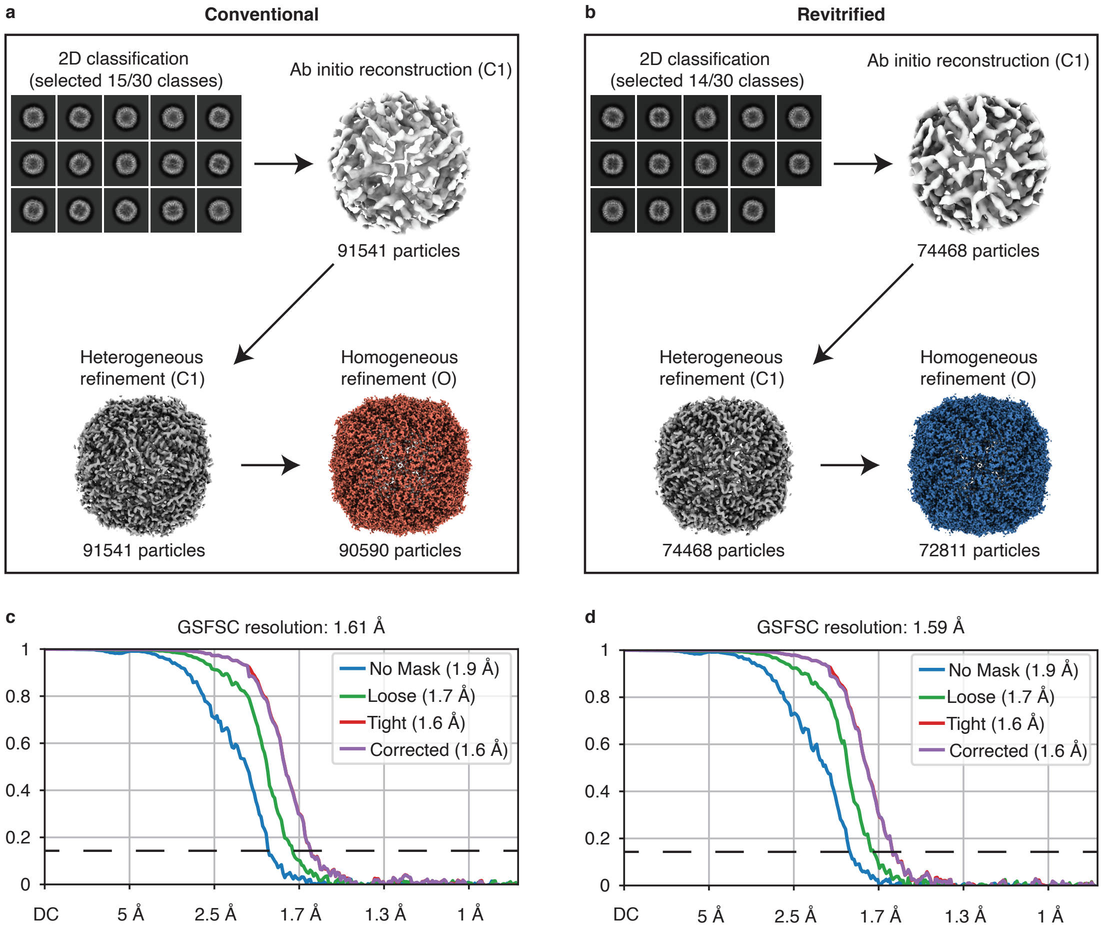


**Supplementary Figure S1. Details of the workflow for single-particle reconstructions from conventional and revitrified sample areas.** (**a,b**) Workflow for the single-particle reconstructions of apoferritin from conventional (**a**) and revitrified (**b**) sample areas. (**c,d**) Gold Standard Fourier Shell Correlations (GSFSCs) for the conventional (**c**) and revitrified (**d**) datasets, with the dashed black line indicating the 0.143 cutoff.
